# ANTARCTIC FUNGI: A BIO-SOURCE ALTERNATIVE TO PRODUCE POLYUNSATURATED FATTY ACIDS (PUFAs)

**DOI:** 10.1101/2022.03.14.484320

**Authors:** Patrizia De Rossi, Alfredo Ambrico, Antonella Del Fiore, Mario Trupo, Luciano Blasi, Marzia Beccaccioli, Luigi Faino, Andrea Ceci, Oriana Maggi, Anna Maria Persiani, Massimo Reverberi

## Abstract

The Antarctic ecosystem is a combination of conditions including extremely low values of temperature. The environmental temperature is one of the parameters thoroughly affecting the structure and composition of fungal membranes lipids. The psychrophilic fungi generally increase the disorder within macromolecules to maintain membrane fluidity at low temperatures. The strategy adopted by Antarctic fungi is to increase the proportion of unsaturated fatty acid that allows maintaining a semi-fluid state of the membranes. This ecological feature might be exploited for using Antarctic fungi as potential alternative source of polyunsaturated fatty acids (PUFAs) for human diet.

This study provides both the characterization of fungal strains isolated from Antarctica by lipidomic analysis and the laboratory/large-scale production of fungal biomass with high content of beneficial PUFAs. In detail, three fungal species isolated from environmental matrices from Antarctica were tested and identified at genome level. Growth experiments to evaluate the influence of temperature and substrate in the yield in biomass and unsaturated fatty acid (UFA) were conducted. The results showed that the selected fungi have a high percentage of UFA compared to saturated ones; low growth temperatures increase the yield in linolenic fatty acid (C18:3); the biomass yield depends on the composition of the growth substrate and a satisfying qualitative-quantitative yield has also been obtained by using an agri-food chain waste product as growth substrate.

**IMPORTANCE:** The presence of polyunsaturated fatty acids (PUFAs) in human and animal diet is gaining attention because PUFAs have several recognized functional properties: they modulate immune response, have anti-allergic and anti-inflammatory activity, cardio-protective effect and reduce blood LDL cholesterol levels. Human diets typically do not contain sufficient PUFAs because foods rich in PUFAs are few and it is therefore necessary to supplement this diet. Food supplements with these types of fatty acid currently commercially available come from marine fish oils and this source is no longer sustainable. It is necessary to develop efficient industrial processes capable of producing good quality PUFAs and in quantity, even using as carbon and nitrogen sources agro-industrial chains’ (in our case spent yeast from brewing and whey waste) waste products. Like microorganisms we used Antarctic fungi because they are adapted at very low temperature increasing the proportion of unsaturated fatty acid that allows maintaining a semi-fluid state of the membranes.

## 1. INTRODUCTION

There is a direct correlation between the adaptation to low temperatures, e.g., in psychrophilic organisms, and the degree of unsaturation of membrane lipids: the lower the temperature, the greater the amount and degree of unsaturation in the fatty acid incorporated into membrane lipids. Some microorganisms, such as fungi, possess a remarkable phenotypic plasticity (1) that allows them adapting, even though epigenetic adaptive memory, to extreme environmental conditions (2, 3, 4). The Antarctic ecosystem is a set of combinations of extreme conditions that include low temperature and humidity, poor availability of organic matter and high salt stress conditions. This leads to lack of diversity and “biomass” contributors to ecological processes (5, 6, 7). The microfungi with their high ratio of surface area/volume, their high phenotypic and metabolic plasticity exist in communities which are simple from a structural point of view but physiologically complex (8).

The interest of this work focused particularly on the phenotypic fungal plasticity in term of polyunsaturated fatty acids (PUFAs) synthesis, as the main adaptive response to low temperatures. PUFAs are found in every organism, but humans cannot synthesize some of them, such as linoleic acid, crucial for the synthesis of others (e.g., arachidonic acid). The presence of polyunsaturated fatty acid, ω3–ω6–ω9, in animal and human diet is gaining attention since PUFAs may act especially as regulators in the blood cholesterol level and as beneficial for the cardio-circulatory system. Among PUFAs, eicosapentaenoic acid (EPA; 20:5n-3) and docosahexaenoic acid (DHA; 22:6n-3), which are n-3 long-chain PUFAs widely referred to as omega-3 oils, were reported to prevent the development of obesity in rodents and humans. In this regard, dietary supplements containing these kinds of fatty acids food supplements with these types of fatty acid are commercially available for health maintenance and the prevention of chronic diseases.

The commercial production of omega-3 oils such as EPA and DHA has primarily depended on marine fish oils to date, but the application of fish oils is often hampered by difficulties including seasonal variations, marine pollution, and the high processing cost (9). In addition, mass-scale fishing to match the increasing demand for fish oils is no longer sustainable (10). The psychrophilic fungi (cold-tolerant), modulating saturated/unsaturated lipids ratio, in response to temperature variations gain recently particular attention as producers of PUFAs, such as gamma-linolenic acid, eicosapentaenoic acid, docosahexaenoic acid, purposed as foods and feeds additives. For an efficient and sustainable microbial PUFAs production, it is crucial to dispose of microbial strains with a naturally high PUFAs content, easily growing in low-cost substrates.

Thus, fungi can be exploited for producing at large scale PUFAs of interest for hum and livestock diets. Nevertheless, the type of substrate used for growing fungi can modify this production. The possibility of producing fungal biomass through fermentation techniques allows controlling a series of growth parameters (pH, dissolved oxygen levels, agitation, nutrients) which leads to a quantitative and qualitative increase of fungal growth and to an increase of production of specific metabolites of commercial interest. Indeed, in recent studies it has also been shown that the composition of the substrate and of the parameters strongly influence the production of PUFAs in microorganisms isolated from seawater and soil (11,12). Furthermore, control of the growth parameters allows standardizing the production of specific compounds and making it repeatable in time.

This study aims seeking for alternative production of PUFAs in a sustainable way by selecting micro fungi isolated in Antarctica growing on substrates deriving from food wastes such as brewery wastes and milk whey.

## 2. MATERIALS AND METHODS

### 2.1 Fungal strains

The following strains were used: *Paecilomyces farinosus*, *Phialophora fastigiata*, isolated from moss of Kay Island (74°05’S, 165°17’E), and a fungal strain of *Agonomycetales* (formerly Mycelia Sterilia) (after identified as *Epicoccum nigrum* – see below), isolated from soil under moss of Starr Nunatak (75°54’S, 162°35E) during 10^th^ summer Italian expedition in Victoria Land Antarctica, 1994-95. The strains were obtained from the culture collection of the Fungi Biodiversity Laboratory (FBL), of the Department of Environmental Biology, Sapienza University of Rome, where they are conserved, respectively, with the numbers FBL 167, FBL 175 and FBL 181.

### 2.2 Growth substrate and cold stress

Preliminary tests were carried out to evaluate the highest biomass yield of the three fungi grown on different liquid media and to evaluate cold stress on fungal grown. In 100 ml flask, 25 ml of sterilized liquid medium with different composition (Tab. 1) were inoculated with 100 μl of fungal suspension (10^4^ CFU/ml) and incubated at 25 °C and 180 rpm. The fungi were cultured to evaluate the growth and their lipidomic profile on different substrate (S10, S11 and S15) at different temperature. In one experiment maintaining the temperature constant at 10°C for 10 days and in another experiment maintaining the temperature for the first 6 days at 25 °C, then lowering it to 10°C for the last 4 days (hereinafter referred to as 25°C&10°C). While, for cold stress trial, flask with only substrate S10 were incubated at 4 °C and 25 °C. After 10 incubation days, the culture liquid was centrifuged at 13000g for 10 minutes at 4 °C. Subsequently the pellet was dried, and the weight of the fungal biomass was determined. All experiments were carried out in triplicate and repeated two times.

### 2.3 Genetic characterization of fungal species

The DNA has been extracted following the 3 C-TAB method (13). Total extracted DNA was visualized by gel electrophoresis. Quantity and quality check was evaluated by spectrophotometer (Nanodrop, Thermo Scientific). 100 ng of pure DNA were amplified by PCR with fungal universal-primers. The couple of universal primers ITS1 (5’-TCCGTAGGTGAACCTGCGG-3’) and ITS4 (5’-TCCTCCGCTTATTGATATGC-3’) amplified the ribosomal DNA (14). Amplification was performed with BIOTAQ™ DNA Polymerase (Bioline, Meridian Bioscience) following the protocol reported by the manufacturer. Identification of fungal species was performed by sequencing the purified amplicon with ISOLATE II PCR and Gel Kit (Bioline, Meridian Bioscience) by Sanger sequencing approach. DNA extraction for Nanopore sequencing was performed as described in Seidl et al. (2015) (15). Nanopore sequencing was performed as manufacture described. The DNA library was construct using SQK-LSK109 and EXP-NBD104 kit and run on a R9.4.1 flowcell sistem.

### 2.4 Production of fungal biomass in a shake-flasks and bench-scale bioreactor

The experiments were conducted in shake-flasks (500 ml) and in a bench-scale volume bioreactor (5 L) using S10, S11 and S15 media. In shake-flasks, 100 mL of medium were inoculated with 100 μl of a fungal suspension at 10^4^ CFU/ml. The fermentations were carried out at 180 rpm and at the different temperature. The bench-scale culture was carried out in a 5-L bioreactor (BIOSTAT B, Braun Biotech International) at working volume of 3 L. For each fungal specie, a preculture of PD broth 96 hours-old was used as starter at rate of 5% (v/v). During the fermentation process the liquid culture was first maintained at pH 7 and temperature of 25 °C for 5 days and then cooled at 10 °C for other 3 days. Aeration and agitation rates were controlled at 2 lpm and 200 rpm, respectively. At end of culturing process, the mycelium was recovered by centrifugation at 13000g for 15 min at 4 °C, washed with sterile water and freeze-dried using a pilot lyophilizer (Christ, Loc-1M). The dried-biomass was gravimetrically determined and stored in vacuum-packaging until to the subsequently lipid analysis. All experiments were carried out in triplicate and repeated two times.

### 2.5 Fatty acid content: extraction and gas chromatography analysis

Fatty acids were extracted from mycelium grown on S10, S11 and S15 medium at 10°C for 10 days and 25°C & 10°C. Fatty acids were derivatized into fatty acid methyl esters following a hydrolysis and methylation-based procedure. To 200 mg of lyophilized mycelia ground in liquid nitrogen were added the nonadecanoic acid internal reference standard for (semi)quantitative analysis and 2 ml KOH 0,5M in methanol; this solution was then maintained at 60°C for 60 minutes. After that, two milliliters of 1 M H_2_SO_4_ in methanol were added and incubated at 60 °C for more 15 minutes. Following the two incubation steps, 2 ml of H_2_O and 2 ml of hexane were added to the tube, vortexed, then allowing the separation of the two lipophilic and hydrophilic phases to take place. The hexane layer containing fatty acid methyl esters (FAME) was removed with a glass pipette and transferred into a gas chromatographic vial. GC was performed on Agilent 7890B GC System, equipped with a FID and a capillary column Omegawax^®^ (30 m 0.25 mm i. d., 0.25 μm film thickness). Injector and detector temperatures were maintained at 250°C and 260°C, respectively. The oven was set at 170°C for 1 min, and then it was increased by one degree per minute (1 °C/min), to 225 °C. The carrier gas, helium, was used at a flow rate of 1.2 ml/min. The injection volume was 1 μl, with a split ratio of 10:1. Methyl esters of palmitic acid, stearic acid, oleic acid, linoleic acid, linolenic acid were used as standard for fatty acid identification and quantification. Two biological replicates are technically repeated four times for the analysis.

### 2.6 Fatty acid content in fungi under cold stress: extraction and gas LC/MS analysis

Fatty acids were extracted from mycelium grown at 4°C, 10°C, 25°C to 10°C (the experiment was carried out maintaining the temperature for the first 6 days at 25 °C, then lowering it to 10°C for the last 4 days))and 25°C on S10 medium for 10 days. 20 mg of lyophilized mycelium were grinded in presence of liquid nitrogen and extracted following the method described in Beccaccioli et al. (16). Internal reference standard was the 9(S)-HODE-d4 (Cayman). HPLC-MS/MS analysis was performed by Single Ion Monitoring (SIM) approach. Fatty acid unsaturation index (UI) indicates the degree of unsaturation in lipids, and therefore the membrane fluidity (17). UI is calculated as the sum of the percentage of each unsaturated FA multiplied by the number of unsaturation present in each FA (18). The formula used is shown below: UI =[1×(% monoenoics)+2×(% dienoics)+3×(% trienoics)]/100. Two biological replicates are technically repeated four times for the analysis.

## 3. RESULTS

### 3.1 Molecular identification of fungal species

The strains (167, 175 and 181) from the Antarctica fungal collection were characterized by sanger sequencing of the internal transcribed spacer (ITS) region (Table S1). BLAST analysis of the three ITS regions showed that the sequence from the isolate 167 match the MN588141.1 representative for *Cordyceps farinosa* clone TR-52-014 (syn. *P. farinosus*), the sequence from the isolate 175 match to *Cadophora fastigiata* strain F-30 (MF077223.1) while the sequence derived from isolate 181 hits to a fungal strain of *Agonomycetales (Epicoccum* type) and shared 98.99% of identity and 99% query coverage with *Epicoccum nigrum* isolate HG12 (KX099630.1). To further strengthen BLAST analysis performed by ITS region, the genome of the three isolates was sequenced by Oxford Nanopore Technology sequencing. DNA sequencing with MinION^®^ generated about 2Gbases for isolates 167 and 175 while 1.6 Gbases for isolate 181 that represents more than 40x coverage for each genome. All three genomes were assembled yielding using mini_asemble software from pomoxis software. The genome size of isolate 167 resulted in 49,641,766 bases in 28 contigs; the gnome size of isolate 175 was estimated in 37,677,608 in 34 while isolate 181 was smaller including 35,337,497 bases over 38 contigs. To assign a taxonomical species, each genome assembly was aligned to about 8500 fungal genomes retrieved from the Assembly NCBI database. The genome of 167 had the best alignment with the sequenced *Cordyceps farinosa* strain KACC 47486 (homotypic synonym

*Paecilomyces farinosus*; GCA_003025275.1) with an identity of about 97.9%, the genome of isolate 181 had the best match with *Epicoccum nigrum* genome from strain P16 with an identity of about 89.5%. Although the identity is quite low, this is the best match that we could get confirming the ITS identification. The alignment between the genome assembly of isolate 175 showed a good identity with *Cadophora malorum* strain M34 with an identity of about 97.7% suggesting that the identification *via* ITS is of a good quality.

### 3.2 Fungal growth evaluation

The effects of several carbon and nitrogen source on biomass are shown in the table 2. For each fungus, the results showed differences in yield which depend on the substrate used for growth. The subsequent tests were conducted with the substrates S10, S11 and S15. These substrates were chosen for the high yield values and S15 also to test the potential of agri-food processing wastes (milk whey and brewery wastes) as low-cost growth substrates for fungi.

**TABLE 1.** Composition of the medium.

| Substrate |  |  | Composition |  |  |  |
| --- | --- | --- | --- | --- | --- | --- |
| S1 | PD broth | 24 g/l |  |  |  |  |
| S5 | Glycerol | 100 g/l | KNO<sub>3</sub> | 10 g/l | Yeast extract | 5 g/l |
| S6 | Glucose | 100 g/l | KNO<sub>3</sub> | 10 g/l | Yeast extract | 5 g/l |
| S7 | Glucose | 100 g/l | KNO<sub>3</sub> | 10 g/l | Corn steep liquor | 5 g/l |
| S10 | Sucrose | 100 g/l | KNO<sub>3</sub> | 10 g/l | Yeast extract | 5 g/l |
| S11 | Maltose | 100 g/l | KNO<sub>3</sub> | 10 g/l | Yeast extract | 5 g/l |
| S12 | Cellulose from paper | 100 g/l | KNO<sub>3</sub> | 10 g/l | Yeast extract | 5 g/l |
| S13 | Potato extract | 100 g/l | KNO<sub>3</sub> | 10 g/l | Yeast extract | 5 g/l |
| S15 | Sucrose | 60 g/l of Mw* | KNO<sub>3</sub> | 10 g/l of Mw* | Brewery wastes | 5 g/l of Mw* |

\*Milk whey

**TABLE 2.**
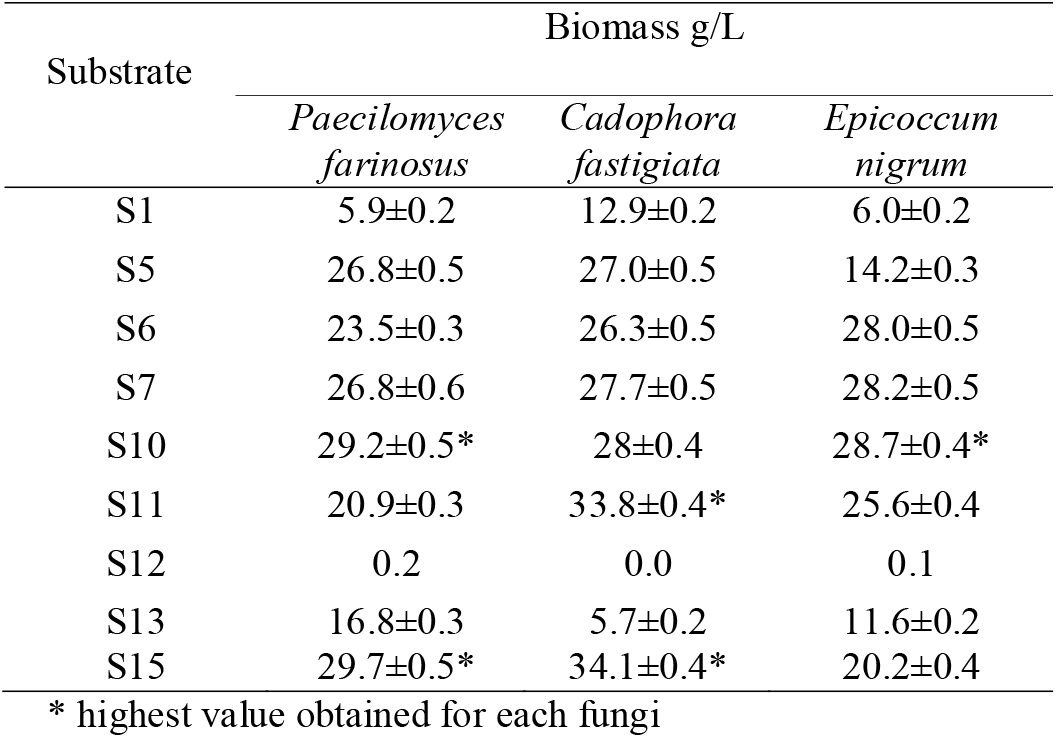
Substrates screening at a temperature of 25 °C

Table 3 shows the production of fungal biomass in a shake-flasks and bench-scale bioreactor expressed in grams of dry biomass per liter of culture medium, at different conditions of substrate and temperature. The results show that yields of biomass in shake-flasks vary from ~20 to 51 g/liter of culture broth and in bench-scale bioreactor vary from ~31 to ~59 g/liter of culture broth. In shake-flasks cultivation, the yields vary from 21 to 38 g/liter of culture for *Paecilomyces farinosus*, from 28 to 51 g/liter of culture broth for *Phialophora fastigiata*, and from 20 to 39 g/liter of culture *Epicoccum nigrum.* The maximum yield in biomass was always obtained at 25°C&10°C, except for *Epicoccum nigrum* in flask at 10°C. In bioreactor the best results in terms of mycelium biomass were achieved with the S15 substrate for all strains.

**TABLE 3.**
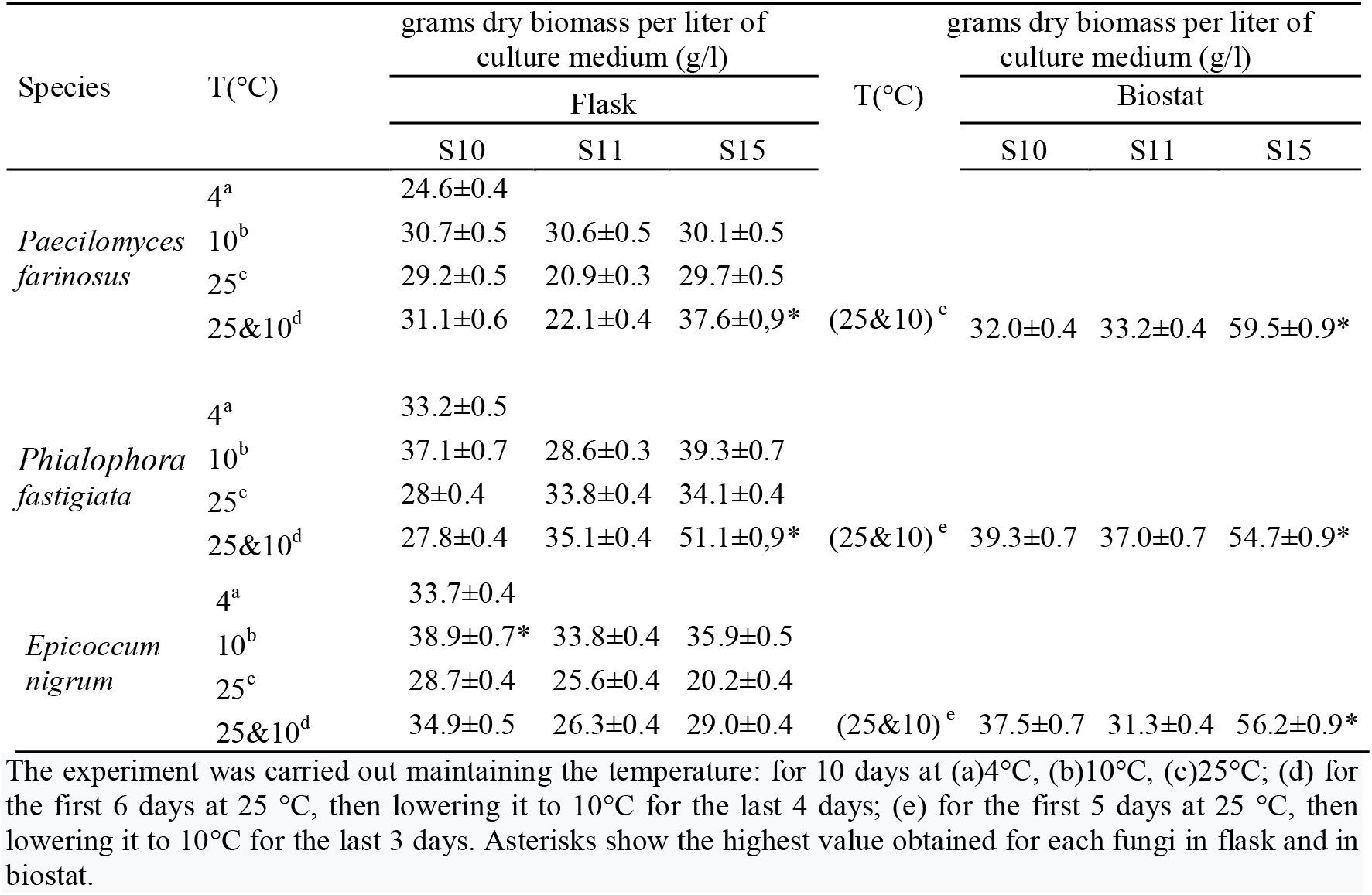
Effect of temperature and medium on the yield of biomass

### 3.3 Determination of fatty acid content in fungi

Here we shown the results of the gas chromatography analysis of *Paecilomyces farinosus, Phialophora fastigiata* and *Epicoccum nigrum* grown on S10, S11 and S15 at 10°C for 10 days and at 25°C&10°C. As shown in figure 1, the total fatty acids (TFAs) contents were included between ~120 and ~570 mg/g mycelia dry weight (dw). In *Paecilomyces farinosus* TFAs contents are between 520 and 570 mg/g mycelia dw the exception was 330 mg/g mycelia dw in substrate S11 at 10°C, value significantly lower (p<0.001). In *Phialophora fastigiata* and *Epicoccum nigrum* we observed a higher total fatty acid content (p<0.001) when the strains were grown at 25°C&10°C compared to growth at 10 °C. The highest values were obtained with the growth on the S15 substrate for *Phialophora fastigiata* (550mg/g mycelia dw) and S11 for *Epicoccum nigrum* (435mg/g mycelia dw). The analysis compared saturated and unsaturated (poly and mono, UFA) fatty acid respectively in flask and in bioreactor (Tables 4A and 4B). The observed proportion of unsaturated fatty acid (UFA) was greater than saturated fatty acid. The highest percentages of UFA were observed with *Epicoccum nigrum* on S10 at 10 °C (~77%). The UFA in *Paecilomyces farinosus* was comprised between ~62% and 65% of the total fatty acid. In *Phialophora fastigiata* UFA amount varied from a minimum of ~55% to a maximum of ~72%. The percentages of UFA in *Epicoccum nigrum* were included between about ~52% and 77%.

**Figure 1.**
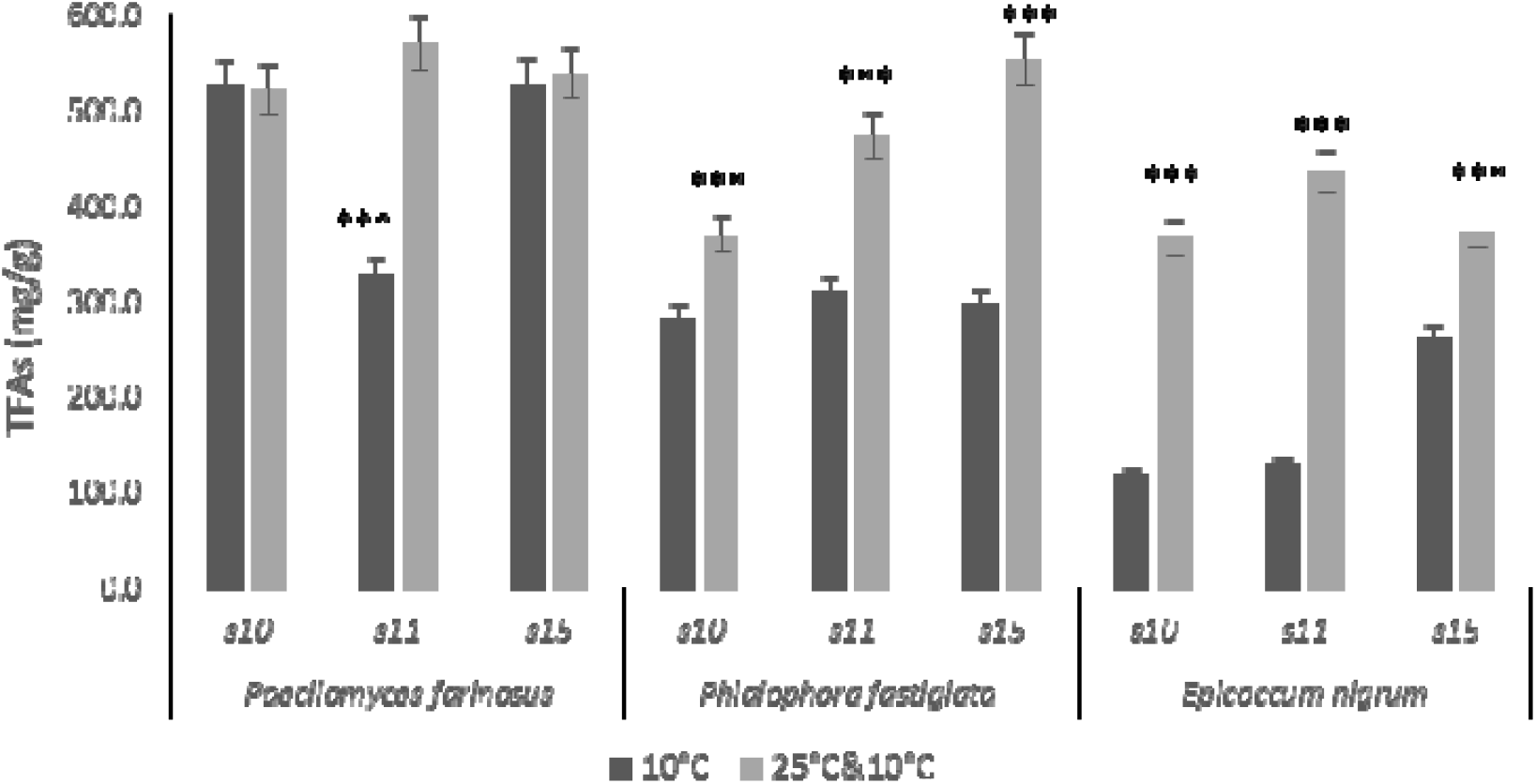
Total Fatty Acids (TFAs) in *Paecilomyces farinosus, Phialophora fastigiata* and *Epicoccum nigrum* grown on different substrate (S10, S11 and S15) at two different temperature conditions:10°C for 10 days and 25° for 6 days then lowering to 10°C for 4 days (25°C&10°C). “g” indicates mycelia dry weight. Results with standard error are means two biological replicates technically repeated four times. “***” indicates *p*<0.001 (strain at 10°C *vs* strain at 25°C&10°C)

**TABLE 4A.** Saturated and unsaturated fatty acids composition in *Paecilomyces farinosus, Phialophora fastigiata* and *Epicoccum nigrum* grown in flakes, at different temperature and culture media

|  |  | Fatty acids production (%) |  |  |  |  |  |
| --- | --- | --- | --- | --- | --- | --- | --- |
| Species | T(°C) | S10 |  | S11 |  | S15 |  |
|  |  | saturated fatty acids | unsaturated fatty acids | saturated fatty acids | unsaturated fatty acids | saturated fatty acids | unsaturated fatty acids |
| *Paecilomyces farinosus* | 10 | 34.4 | 65.6 | 34.2 | 65.8 | 37.5 | 62.5 |
|  | 25&10 | 36.4 | 63.4 | 35.2 | 64.8 | 36.7 | 63.3 |
| *Phialophora fastigiata* | 10 | 37.5 | 62.5 | 27.6 | 72.4 | 45.1 | 54.9 |
|  | 25&10 | 31.8 | 68.2 | 27.6 | 72.4 | 38.2 | 61.8 |
| *Epicoccum nigrum* | 10 | 23.3 | 76.7 | 30.8 | 69.2 | 48.0 | 52.0 |
|  | 25&10 | 30.0 | 70.0 | 25.5 | 74.5 | 33.9 | 66.1 |

Fungi grown on S10, S11, S15 at 10°C for 10 days and at 25° for 6 days then lowering to 10°C for 4 days (25°C&10°C). Values expressed as percentage of the total fatty acid concentration

**TABLE 4B.** Saturated and unsaturated fatty acids composition in *Paecilomyces farinosus*, *Phialophora fastigiata* and *Epicoccum nigrum* grown in bioreactor on different culture media

| Species | Fatty acids production (%) |  |  |  |  |  |
| --- | --- | --- | --- | --- | --- | --- |
|  | S10 |  | S11 |  | S15 |  |
|  | saturated fatty acids | unsaturated fatty acids | saturated fatty acids | unsaturated fatty acids | saturated fatty acids | unsaturated fatty acids |
| *Paecilomyces farinosus* | 36.0 | 64.0 | 34.3 | 65.7 | 35.0 | 65.0 |
| *Phialophora fastigiata* | 20.3 | 79.7 | 29.3 | 70.7 | 31.7 | 68.3 |
| *Epicoccum nigrum* | 31.8 | 68.2 | 36.8 | 63.2 | 33.2 | 66.8 |

Fungi grown on S10, S11, S15 at temperature of 25 °C for 5 days and then cooled at 10 °C for other 3 days (25°C&10°C). Values expressed as percentage of the total fatty acid concentration.

The table 5 shows the fatty acid quali-quantitative profile of the fungal strains: C16:0 palmitic (14-36%), C18:0 stearic (7-19%), C18:1 oleic (16-48%), C18:2 linoleic (16-41%), C18:3 linolenic (0.2-25%) acids. These fatty acids were present in all the fungal strains analyzed. The most abundant fatty acids were palmitic, oleic and linoleic acid, representing ~70-90% of the total fatty acid analyzed. The percentages of linolenic acid (18:3) were highest at 10°C, lowest values were observed at 25°C&10°C regardless of the substrate used. *Phialophora fastigiata* showed the highest percentages of linolenic acid in all the growth conditions tested.

**TABLE 5.** Palmitic, stearic, oleic, linoleic, linolenic acids composition in Antarctic fungal species at different condition and culture media

| Species | T(°C) | substrate S10 |  |  |  |  | substrate S11 |  |  |  |  | substrate S15 |  |  |  |  |
| --- | --- | --- | --- | --- | --- | --- | --- | --- | --- | --- | --- | --- | --- | --- | --- | --- |
|  |  | C16:0 | C18:0 | C18:1 | C18:2 | C 18:3 | C16:0 | C18:0 | C18:1 | C18:2 | C 18:3 | C16:0 | C18:0 | C18:1 | C18:2 | C18:3 |
| *Paecilomyces farinosus* | 10 | 25.0 | 9.4 | 34.4 | 28.8 | 2.4*** | 23.3 | 10.9 | 27.6 | 34.8 | 3.4*** | 21.3 | 16.2 | 34.2 | 25.9 | 2.4*** |
|  | 25&10 | 27.8 | 8.8 | 33.7 | 28.4 | 1.3 | 24.1 | 11.1 | 36.0 | 27.7 | 1.1 | 23.48 | 13.2 | 39.4 | 22.6 | 1.3 |
| *Phialophora fastigiata* | 10 | 20.6 | 16.9 | 27.7 | 19.8 | 15.0*** | 19 | 8.6 | 24.8 | 31.4 | 16.2*** | 26.4 | 18.7 | 19.2 | 16.5 | 19.2*** |
|  | 25&10 | 23.4 | 8.4 | 16.4 | 38.3 | 13.5 | 19.7 | 7.9 | 26.8 | 33.7 | 11.9 | 24 | 14.2 | 26.3 | 25.7 | 9.8 |
| *Epicoccum nigrum* | 10 | 13.6 | 9.7 | 41.6 | 32.2 | 2.9*** | 17.8 | 13.0 | 29.5 | 37.0 | 2.7*** | 34.1 | 13.9 | 25.9 | 23.5 | 2.6*** |
|  | 25&10 | 15.7 | 14.3 | 37 | 31.9 | 1.1 | 16.7 | 8.8 | 42.4 | 31.5 | 0.6 | 24.4 | 9.5 | 34.9 | 30.6 | 0.6 |

Fungi grown on S10, S11, S15 at 10°C for 10 days and at 25° for 6 days then lowering to 10°C for 4 days (25°C&10°C). Values expressed as a percentage of the total fatty acids concentration. “\*\*\*” indicates $p < 0.001$ (strain at 10°C vs strain at 25°C&10°C). In bold, showed the highest percentages of linolenic acid (18:3) obtained for each fungi.

To verify whether the differences between the concentrations of linolenic acid detected at variable temperatures were significant, a statistical analysis was carried out. The result showed percentages in linolenic acid always statistically significant (p<0.001).

### 3.4 Fatty acid composition in *Paecilomyces farinosus, Phialophora fastigiate* and *Epicoccum nigrum* under cold stress condition

Synthesis and composition of FAs determine the fluidity of membranes. We performed FA analysis of *Paecilomyces farinosus* (#167), *Phialophora fastigiata* (#175) and *Epicoccum nigrum* (#181) grown on S10 at 4°C and 25°C. Saturated and unsaturated FAs were considered. In general, we observed a significant higher level of total fatty acid when strains were grown at 25°C, suggesting the influence of the temperature on the FA synthesis (Fig. 2). To describe the incidence of unsaturated FAs and their role on the membrane composition and fluidity we introduced the Unsaturation Index (UI). The UI represents the concentration of the unsaturated FAs, a low index corresponds to a lower amount of unsaturated FAs so a reduced fluidity. Results in Fig.2b showed that the UI is lower in each strain grown at 25°C, suggesting the increase of the unsaturation is an adaptive condition of these fungi.

**Figure 2.**
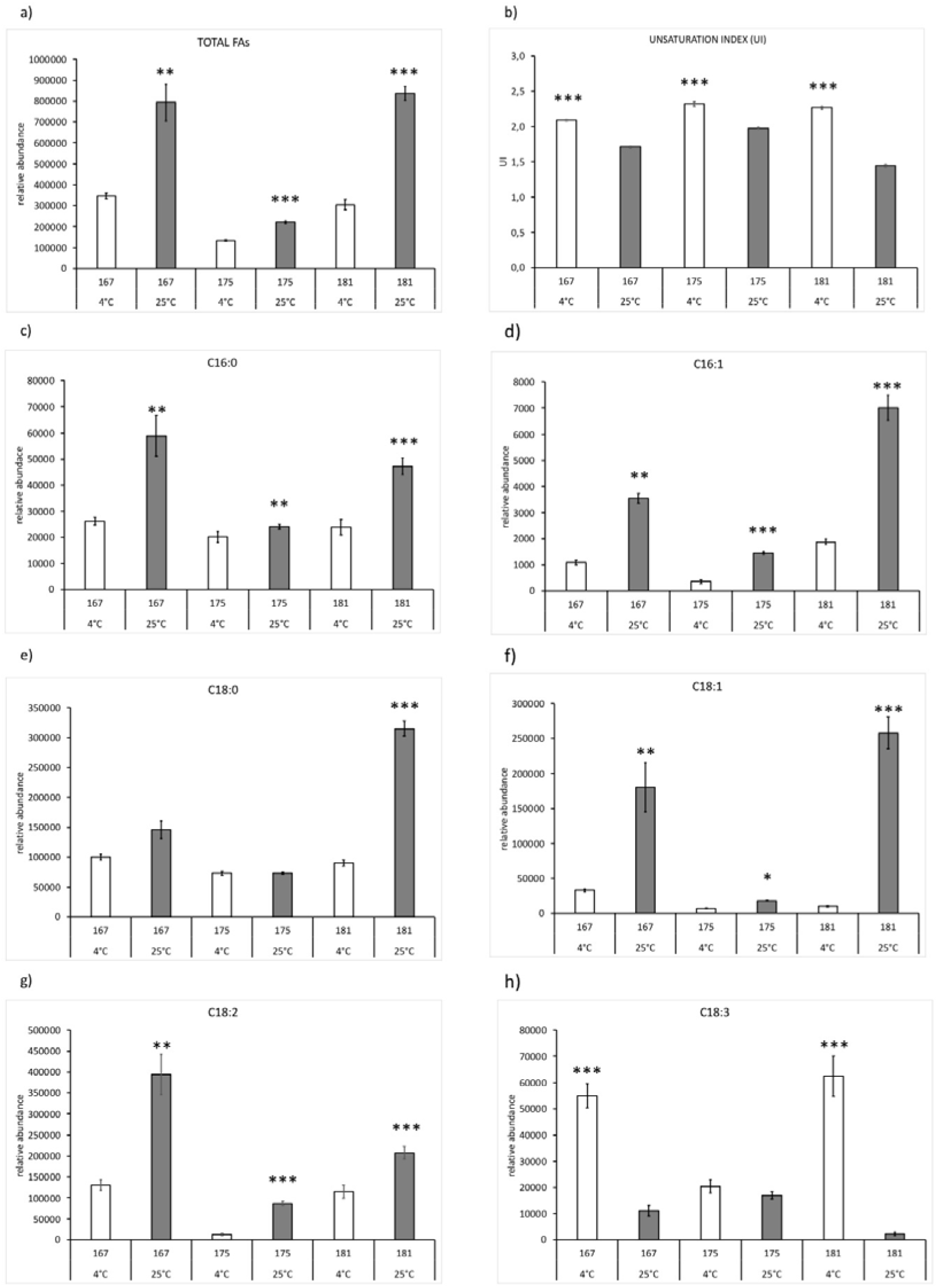
Fatty acid composition in *Paecilomyces farinosus* (#167), *Phialophora fastigiata* (#175) and *Epicoccum nigrum* (#181) grown on S10 at 4°C and 25°C for 10 days. Panel a) report the total FAs; in b) Unsaturation Index (UI) is reported. Panels c-h represent the relative abundance (Area FA/Area internal standard 9-HODEd4) of several FAs. Results with standard error are means two biological replicates technically repeated four times. Asterisks show the statistical relevance: strain at 4°C *vs* strain at 25°C. * *p*<0.05, \*\**p*<.0.01 and \*\*\**p*<0.001.

Palmitic (C16:0), palmitoleic (C16:1), stearic (C18:0), oleic (C18:1), linoleic (C18:2) and linolenic (C18:3) acids have been analysed. The growth at 25°C provokes an increase of the saturated palmitic acid in all the three strains (Fig. 2c), while the stearic acid result more abundant only in the *Epicoccum nigrum* (#181) (Fig. 2e). Palmitoleic and oleic monounsaturated FAs are induced in all the three strains growth at 25°C (Fig.2 d, f). The linoleic acid, a polyunsaturated FA, result more abundant in the strains grown at 25°C (Fig. 2g) but the linolenic acid, with three degrees of unsaturation, is most represented in the strains grown at 4°C (Fig. 2h). In fact, it already been reported that the low temperature induces the increase of the proportion of unsaturated fatty acid (19, 20). In particular, our growth experiments conducted at different temperatures (25°C, 25°C&10°C, 10°C and 4°C), fig. 3, show that lowering the growth temperature from 10°C to 4°C the induction of linolenic acid increases more than it increases from 25°C to 10°C. In fact, in *Paecilomyces farinosus* the percentage of linolenic acid, at 10°C, is 2.4% and becomes 15.9% at 4°C, in *Epicoccum nigrum* it is respectively 2.9% and 20.5%. In *Phialophora fastigiata*, on the other hand, the maximum induction of linolenic acid (about 15%) is reached already at 10°C.

**Figure 3.**
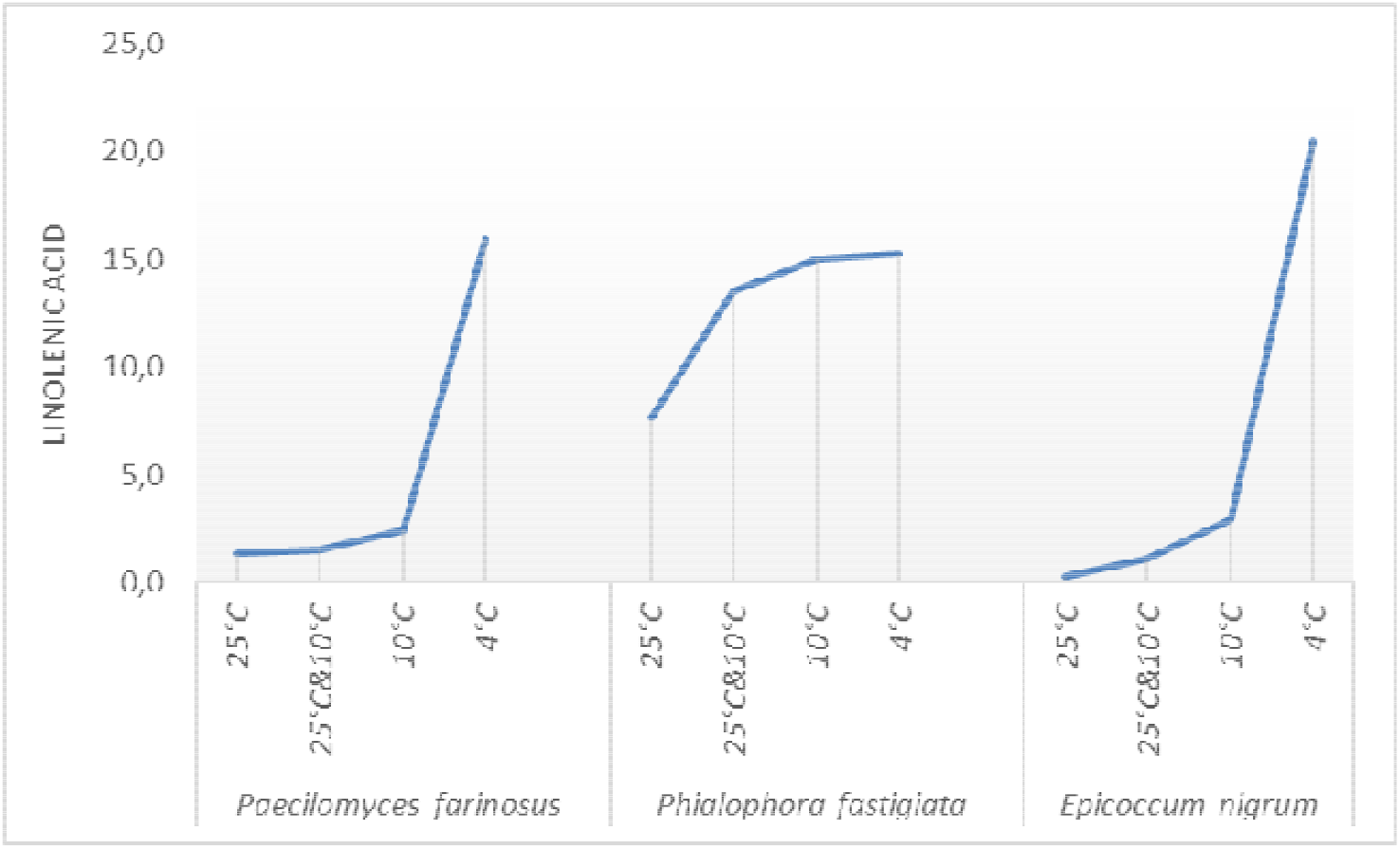
Linolenic acid (expressed as a percentage of the total fatty acids concentration) in *Paecilomyces farinosus, Phialophora fastigiata* and *Epicoccum nigrum* grown on S10 at 25°C, 25°C&10°C, 10°C and 4°C for 10 days.

## DISCUSSION

Aware that microorganisms adapted to live in permanently cold environments represent an unexplored source of potentially applicative biodiversity, this study wanted to recognize, among those, new species to be used to develop competitive processes to produce microbial oils. The idea of identifying oleaginous species among those adapted to live in habitats with temperatures close to zero was plausible since the essential role of highly polyunsaturated fatty acid stored in membrane phospholipids to maintain membrane functionalities in environments permanently cold is known. Changes in lipid metabolism are the major responses through which acclimatization and adaptation to cold in microorganisms, in general, are implemented; this is explained by the need to maintain the appropriate physiological state and therefore the functionality of the lipid membranes, otherwise compromised at temperatures close to freezing (21,22).

Different microorganism types can synthesize long-chain polyunsaturated fatty acids, among which there are bacteria, yeasts, filamentous fungi, and microalgae (23, 24). The extraction of oils from microorganisms, have some advantages, such as shorter life cycle and abundant and cheap raw material, easy to be produced on a large scale (25). The fungi, moreover, can grow up under different conditions, with consequent different production of lipids on the different substrates (26). Therefore, is important to identify parameters to optimize polyunsaturated fatty acid production (27,28).

In the present work, for all studied fungi we obtained high yields of mycelial dry matter in many of the substrates tested in preliminary screening (tab 2) if compared with those obtained in PDbroth (substarte S1). In particular, these latter are ranging from 5.9 to 12.9 g/L. and are in according with yields reported in other independent reports. For example, Liu *et al.* and Mascarin *et al.* (29, 30) culturing *Isaria farinose* in shake-flasks with PDbroth at 25 °C reported yields of 5.83 and 4.28 g/l, respectively. While Salehi *et al.* (31) for *Epicoccum nigrum* have achieved values of biomass between 3-12 g/L on Murashige and Skoog medium. The highest yields obtained with other substrates tested by us, exception make for S12, could be due to the high concentration of the carbon sources used in the media. This claim is supported by the findings observed for other fungi species in several studies. Zhu *et al.* (32) in shake-flasks obtained 31.2 g/L of dry-biomass culturing *Mortierella alpina* on inexpensive medium composed by maize starch hydrolysate (with glucose concentration of 100 g/L) and different nitrogen source. Kim *et al.* (33) reported that in shake-flask cultures the mycelial growth of *Paecilomyces sinclairii* was proportionally to the increasing of sucrose concentration within the ranges from 10 to 60 g of sugar/L.

Furthermore, in flasks we observed that the most interesting experimental conditions for fungal biomass production were 25°C&10°C, except for *Epicoccum nigrum* 10°C, S10 and S15. The maximum yield in biomass is obtained in bench-scale bioreactor for *Paecilomyces farinosus* on media S15. The condition 25°C&10°C allowed to obtain a higher content of fatty acids than the condition at 10°C observing a total fatty acids content up to about 570 mg/g of dry cell weight. Although under different conditions, Ho and Chen (34) obtained a total fatty acids content in *Mortierella alpina* up to about 400 mg/g of dry cell weight. The proportion of unsaturated fatty acid (UFA) was greater than saturated fatty acid with percentages ranging between 62 to 77%. Lowest values in shake-flasks using the S15 substrate, were found. This could be explained by the presence of whey waste in the S15 medium. In fact, it was observed that cultures in an acid medium (as whey waste is) showed an increased synthesis of saturated fatty acid that enhance the rigidity of the cell membrane to maintain proper cell function during stress conditions (35). Also, in our case the higher percentages of UFA obtained in the Bioreactor could be attributed to the automatic control of acidity (pH 7.0) during the fungal growth. Unlike what has been observed so far, the percentages of linolenic acid (18:3), a precursor of fatty acid of great interest for human health, were higher when the fungi grew at 10°C compared to growth at 25°C&10°C. Maggi et al, 2013 (36) observed that the cold experimental condition (10°C) determines a higher concentration of linolenic acid with respect to a growth at 25°C. The results obtained by Xian M. et al 2001 (37) also showed that the proportion of linolenic acid increased with a decrease in the cultivation temperature and reached a maximum at 5°C. We tested these experimental conditions (i.e. growing the fungi at 5°C) obtaining similar results: the percentage of unsaturation level increased significantly in the cold stress-treated samples thanks to the rise of the level of linoleic acid at 5°C. This increase has been explained by the role of PUFAs in enhancing the membrane fluidity. At lower temperatures, more oxygen is dissolved in water and, thus, more oxygen is available for the oxygen dependent desaturase enzymes (38, 39).

In our results, palmitic (16:0), stearic (18:0), oleic (18:1), linoleic (18:2) and linolenic (18:3) acids were present in every fungal strains. These results are like the findings by Stahl & Klug (40), who determined that the aforementioned fatty acids were the most common and abundant after analyzing 100 strains of filamentous fungi, including Oomycetes, Zygomycetes, Ascomycetes, Basidiomycetes, and *Agonomycetales* (*formerly Micelia sterilia*). In all the samples it was observed that palmitic, oleic and linoleic fatty acids in the fungal mass are mainly represented, confirming the literature data (41).

Our results suggest that the examined fungi could represent an interesting bio-source to produce PUFAs, both in qualitative and quantitative terms. Using agro-food processing wastes growth media increase the competitiveness of oleaginous organisms against conventional PUFAs producers.

